# An integrative assessment of aquatic ecosystem services based on guideline thresholds

**DOI:** 10.1101/768200

**Authors:** Nicolas F. S-Gelais, Jean-François Lapierre, Robert Siron, Roxane Maranger

**Affiliations:** Département des Sciences Biologiques, Université de Montréal, Montréal, Québec, Canada; Groupe de Recherche Interuniversitaire en Limnologie (GRIL); Ouranos–Consortium on Regional Climatology and Adaptation to Climate Change, Montréal, Québec, Canada

## Abstract

Ecologists typically associate water quality with trophic status where oligotrophic ecosystems have excellent water quality and presumably provide more aquatic ecosystem services. However water quality is perceived differently among worldviews. Aquatic ecosystem service provisioning to the public health and agriculture sectors is determined using specific guidelines. But are these guidelines related to trophic status? Here, we developed an integrative ecosystem service framework using guideline thresholds for drinking, swimming, irrigation, suitability for livestock and aquatic wildlife in canadian rivers of varying trophic status. Drinkability was the most sensitive ecosystem service, met in 37% of cases, whereas livestock was the least, provided in 99%. Trophic status is a fair proxy for ecosystem services limited by fecal contamination as nutrients are related to human and animal populations, but not to those limited by metals. Using quantitative thresholds to assess the safe provisioning of multiple ecosystem services provides clear guidance for supporting resource management.

**In a nutshell:**

- Water quality is a commonly used term in management, but the metrics that determine whether a river can safely provide various aquatic ecosystem services differ among worldviews.
- We propose an integrative approach based on guideline thresholds to evaluate the frequency with which rivers are drinkable, swimmable, suitable for irrigation, livestock, and aquatic wildlife and compared this suitability with trophic status.
- Trophic status is a fair proxy for ecosystem services limited by fecal contamination, but not for those limited by metals.
- Using and developing more guideline thresholds provides a concrete way to assess ecosystem service provisioning that could help serve management.

## 1. Introduction

### 1.1 Challenge

Water quality means different things to different people; but regardless of your worldview, an aquatic ecosystem with good water quality supposes that it is capable of providing multiple aquatic ecosystem services (UNEP 2016). Guideline thresholds of specific water usages provide a means to evaluate whether multiple aquatic ecosystem services can be delivered simultaneously or not. This could serve as a novel and valuable approach for management, but has never been applied in an ecosystem service context to our knowledge. From an ecological worldview, trophic status is known to directly and indirectly affect the delivery of multiple ecosystem services. For example, excess nutrients, as a function of human activities on the landscape, can result in toxic cyanobacterial blooms, anoxic conditions, and biodiversity loss resulting in significant environmental and economic damages (Orth et al. 2006, Dodds et al. 2006, Carpenter et al. 2011). But how well does trophic status serve as a proxy to the delivery of multiple aquatic ecosystem services that are typically assessed using guideline thresholds? Using data from Canadian rivers, we propose a novel integrative aquatic ecosystem service framework that tests the hypothesis of whether trophic status is an overarching indicator of water quality.

### 1.2 Water quality across worldviews

Water quality can be defined in multiple ways depending on one’s worldview. In order to explore these potential differences, we compared the use of the term *water quality* throughout the literature across the following research fields: ecological, public health, agricultural, and biodiversity sciences. The use of the term *water quality* has gained traction within the scientific literature across these four fields since the late eighties (Figure 1). This acceleration may be expected given the major amendments to water acts around that period in many different countries including the United States, Canada and the United Kingdom (Canada Fisheries Act 1985, US water quality act 1987, CEPA 1988, UK Water act 1989). But are the reasons why we observe this trend similar among fields? In order to evaluate this, the next step was to compare the most commonly used words around water quality with respect to each worldview.

**Figure 1:**
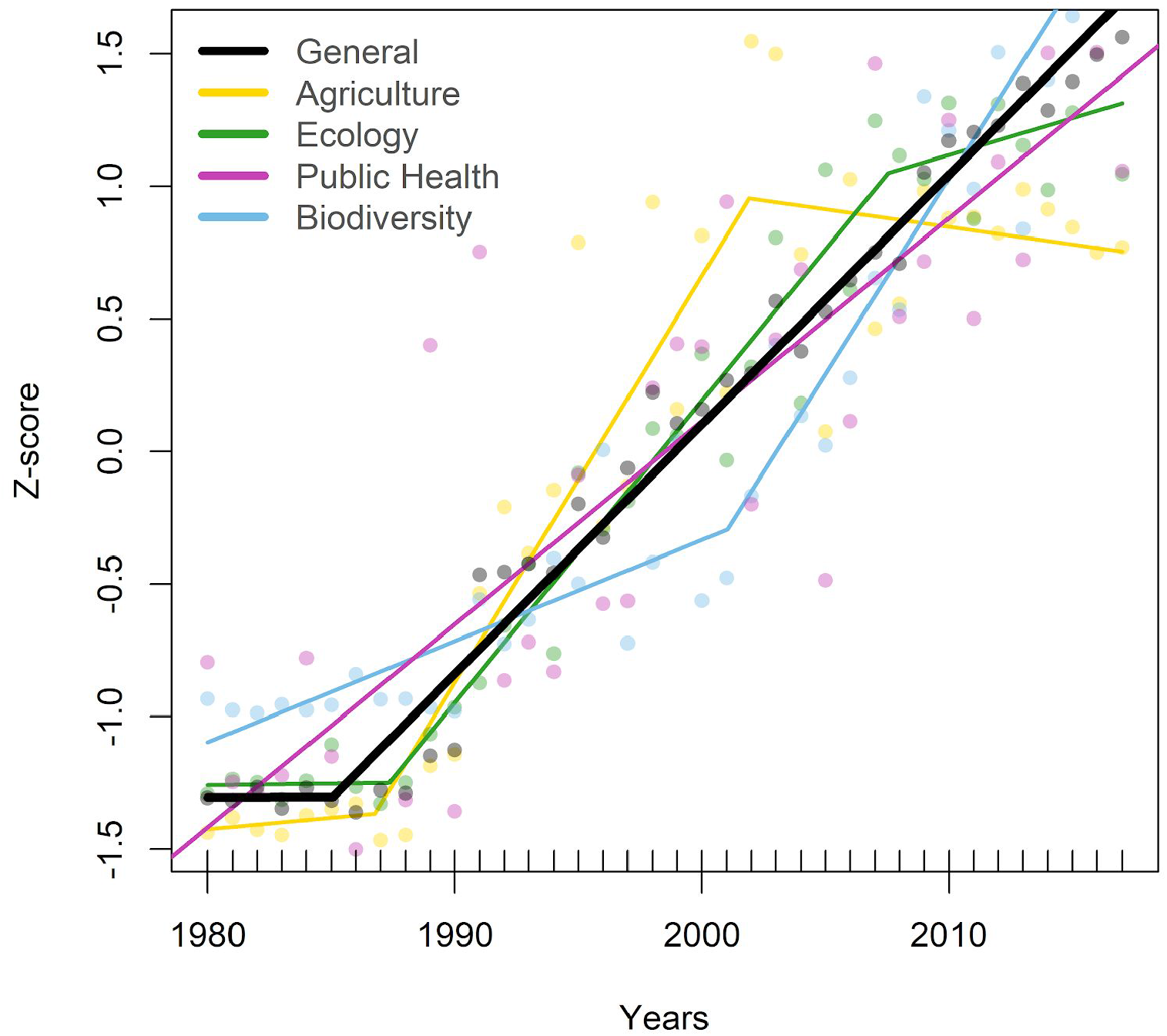
The temporal evolution (from 1980 to 2017) of the relative proportion (Z-score) of scientific papers with the term water quality in the title or abstract in the scientific literature in the fields of ecology, biodiversity, agriculture and public health, based on Web Of Knowledge Core Database. See Table S1 and WebPanel 1 for details.

We extracted the words most often associated with *water quality* in abstracts from over 600 articles published in 2017 in the four fields (Table S2). We then measured the cosine distance between each field (WebPanel1, Table S3) to compare the co-occurrence of words. We found that the language used around public health was the most distinctive of the worldviews compared. Ecology was more similar to both agriculture and biodiversity, but since there was little overlap between the latter two, the reasons for association with ecology were different. In particular, we observed that in ecology 45% of abstracts included at least one of the following words: *nutrient(s)*, *phosphorus,* or *nitrogen*. This is consistent with the notion ecologists have of trophic status providing an indication of aquatic ecosystem health through low algal biomass and water clarity (Anders and Ashley 2007). By comparison, in public health, within the top 10 words were *drinking, risk, contamination, and E. coli* (Table S2). This emphasizes the worldview around the main water use of public concern, clean drinking water, and the metric of contamination most often used to evaluate potability is the presence of *Escherichia coli* (E. coli, Table S2). For agriculture, water is withdrawn for crop irrigation or livestock production which both have different criteria to assess suitability for each aquatic ecosystem services. However, agriculture itself impacts water quality and is globally recognized as the main source of nutrient pollution to aquatic ecosystems (Gordon et al. 2010, Mateo-Sagasta et al. 2017). This is reflected in the keywords which include components of the different ecosystem services for agriculture (e.g. crop yield, irrigation) and words associated with nutrients including *nitrogen*, *phosphorus*, and *nitrate*, which were present in 51% of the abstracts evaluated. In the biodiversity literature, the words associated with water quality were more similar to what we observed in ecology, although nutrients did not emerge while the more frequently used terms were related to diversity metrics (e.g. *abundance*, *index*, *indicators*).

Overall there is surprisingly little overlap in the terms associated with water quality among worldviews. One possible explanation for this divergence is that every worldview focuses on different aquatic ecosystem services each with their own threat to provisioning that serves as the guideline threshold. One of the few exceptions was the term *management*, which was used across all fields. This emphasizes the need to manage ecosystems to maintain water quality for multiple ecosystem services, which integrates different worldviews.

## 2. Aquatic ecosystem service guidelines among worldviews

### 2.1 Threshold guidelines of different aquatic ecosystem services

To assess the provisioning of multiple ecosystem services at once while evaluating their potential tradeoffs, an ecosystem service bundle approach has been proposed (Raudsepp-Hearne 2010). This, however, is typically done using continuous variables whereas much of water quality management often relies on guidelines using thresholds (Hart 1993). This offers an easy to implement, binary decision-making tool on whether an ecosystem service can be delivered or not, bridging management with ecosystem service science (Daily et al. 2009, Guerry et al. 2015). Given that many aquatic ecosystem services have their own set of biological and/or chemical guidelines, we assessed the ability of an aquatic ecosystem to safely provide conditions for drinking, irrigation, swimming, livestock, and biodiversity maintenance simultaneously. Thus, we were able to test several provisioning ecosystem services, a cultural one as well as biodiversity maintenance which is considered by some a regulatory ecosystem service and by others a regulator of ecosystem processes (Mace et al. 2012). In fact it can even be considered more broadly within the context of recreation and provisioning with regards to fisheries for example (Ziegler et al. 2017).

Guidelines for water quality assessments across worldviews are typically divided into five different classes that include biological indicators, inorganic chemicals, heavy metals, pesticides, and other organic pollutants (Health Canada 2017). Although water quality guidelines are regionally distinctive (Boyd 2006), for this exercise we used a series of Canadian guidelines. They were selected for their measurement frequency in publicly available databases as well as for their use in assessing multiple ecosystem services (Table S4). For livestock and irrigation, guidelines were derived from the Canadian Council of Ministers of the Environment (CCME 1999), whereas for drinking and recreation, these came from Health Canada (Health Canada 2012, 2017). Finally, for biodiversity, we used the CCME guidelines for the protection of aquatic life (CCME 1999), which we considered as a proxy to the threat to biodiversity maintenance in a given aquatic ecosystem. For the ecological worldview, we used the trophic level criteria established by Dodds (1998) for rivers that are based on specific nutrients and chlorophyll *a* concentrations.

### 2.2 Overlap between public health, agriculture, and biodiversity guidelines; ecology stands alone

We observed three distinctive ecosystem service clusters (Table S5), which were largely based on the commonality of the metrics used for the guideline thresholds across fields. Drinking, livestock, irrigation, and the protection of aquatic wildlife all cluster quite strongly since they are all based primarily on the same, multiple toxicological guidelines (metals, pesticides, and other organic chemicals; Table S4). Although drinkability and irrigation cluster with recreation, the result is much weaker. This is likely due to the fact that they all have guidelines associated to fecal contamination, but contrary to drinkability and irrigation, which also have toxicological guidelines, recreation is based on fecal contamination alone.

The ecological worldview is the most distinctive as total phosphorus and total nitrogen are not used as guidelines to evaluate any of the other ecosystem services. While there is little debate among ecologists that ecosystems tend to degrade as they become more eutrophic (Smith 2003), how this degradation impinges on the safe delivery of aquatic ecosystems services based on guideline thresholds remains unknown. Therefore, we aimed to empirically test the widely assumed, but often implicit notion that ecosystems with lower trophic status have better water quality in terms of their capacity to deliver multiple aquatic ecosystem services.

## 3. Trophic status as a proxy of aquatic ecosystem service delivery

### 3.1 A multiple ecosystem service framework based on guideline thresholds

We aimed to test the assumption that the frequency with which ecosystem services are provided decreases from oligotrophic to mesotrophic to eutrophic rivers. As a first step, we quantified the overall frequency with which Canadian rivers were able to deliver multiple ecosystem services. Secondly, we evaluated how well trophic status allowed us to predict the capacity of rivers to deliver these ecosystem services. To do so, we used open water quality data from 46 datasets on rivers across Canada (see WebPanel 2). We grouped data by sampling event, which we defined as water collection at a given location in a given month for a total of over 60 thousand observations. However, there were many gaps in this collated dataset. Although over 96% had available information on trophic status based on total phosphorus concentrations, information was more sparsely available for variables used as guidelines. There were 56 % on average for different metals, 1% on average for organic pollutants, 55% for inorganic pollutants and only 11% for E. coli (Table S6). The ability to deliver an ecosystem service was evaluated on the basis of each available guideline for a given sampling event. We focused on the 11% of observations that measured E. coli because it is a major limiting factor for drinkability, and irrigation, and the only guideline for swimmability (Table S6).

### 3.2 What limits the delivery of different ecosystems services across rivers

We used a radar plot to visualize the total frequency with which rivers could provide different ecosystem services based on thresholds (Figure 2a). Overall, the suitability of a river to serve as a source of drinking water for livestock was by far the least sensitive ecosystem service evaluated, where guidelines were not met in only 3 events (Figure 2a). In the rare cases where water wasn’t suitable for livestock, it was limited by high concentrations of heavy metals such as cadmium and lead. Aquatic wildlife was also mostly limited by cadmium and lead, however, the suitability of water to protect biodiversity was much lower at 42% of sampling events. This result should be interpreted with caution because although we assumed the guidelines for the protection of aquatic wildlife could serve as a proxy for biodiversity maintenance, the thresholds are based on the response of the most sensitive species of a predefined multiple-proxy toxicological dataset (CCME 2003, Table S7). Therefore suitability for aquatic wildlife should be placed in a site-specific context to correct for natural background concentrations and species composition.

**Figure 2:**
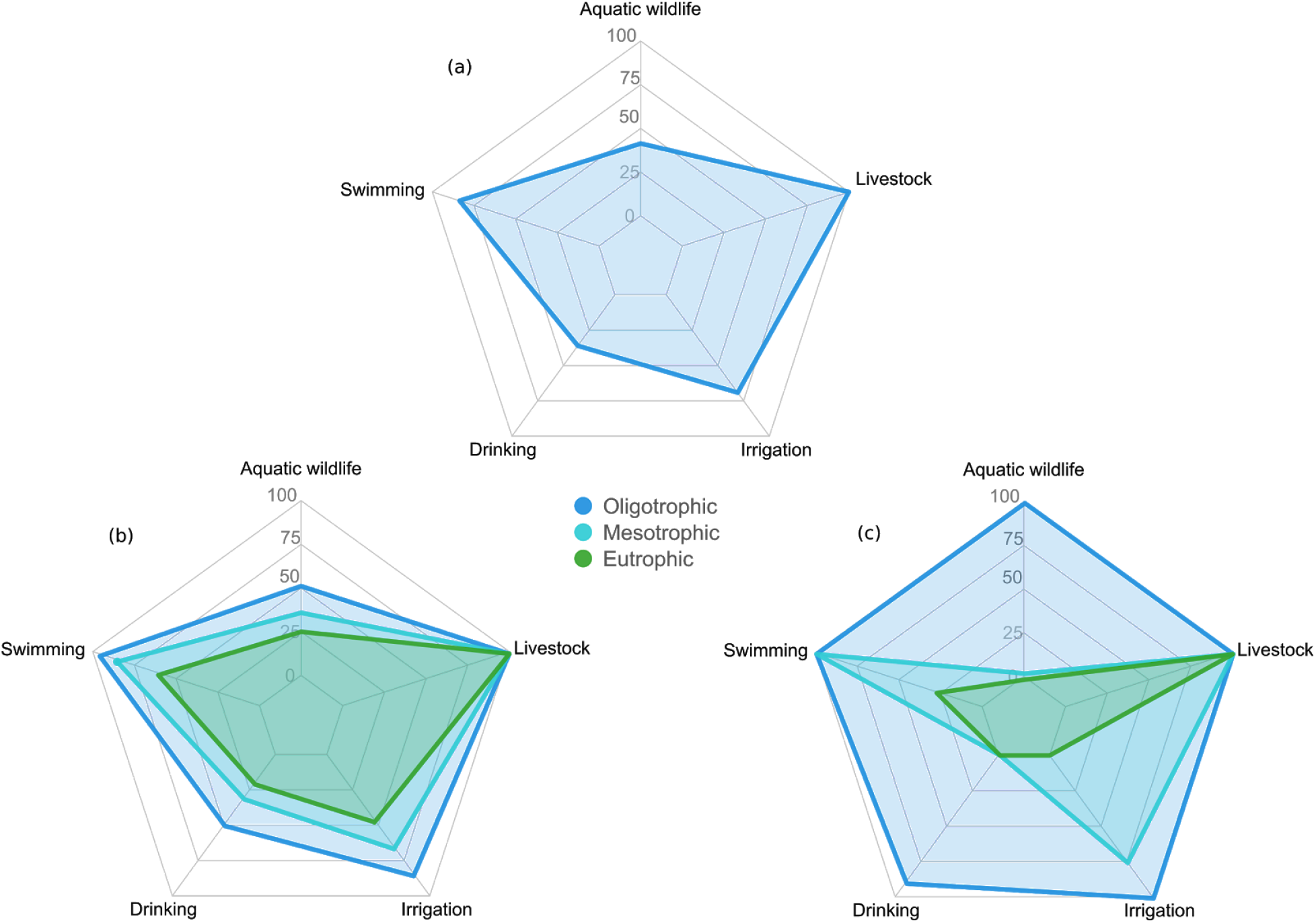
Radar plots of the proportion of sampling events for which aquatic ecosystem services could be provided in (a) all sampling events; (b) in oligotrophic, mesotrophic and eutrophic systems; and (c) what would be expected if trophic status is a perfect proxy of whether or not a service can be provided.

By comparison the suitability for drinking, irrigation, and recreation was almost always limited by E. coli (Table S6). Recreation was always limited by E. coli as it was the only metric of evaluation, whereas heavy metals limited drinking and irrigation in 2 % of events; neither of these services were impinged by pesticide concentrations. Rivers were swimmable in 84% of sampling events whereas irrigation and drinking were safe in 70% and 37% of cases, respectively. Drinking is clearly the most sensitive ecosystem service, since drinking water should be exempt of any fecal coliform.

### 3.3 Trophic status as a proxy of aquatic ecosystem service provisioning

We evaluated the adequacy of trophic status as a proxy for ecosystem services by comparing the proportion of oligotrophic, mesotrophic, and eutrophic ecosystems that met Canadian guidelines (Figure 2b). Overall a larger proportion of ecosystem services were rendered in more oligotrophic ecosystems (mean of 82% across services), whereas fewer were rendered in mesotrophic (mean of 62% across services) and eutrophic ones (mean of 46% across services), confirming the hypothesis that trophic status can inform on ecosystem service provisioning.

Not all ecosystem services were equally compromised by trophic status. Drinking, swimmability, and irrigation were the most responsive to trophy, as they tended to be limited by E. coli, which is generally well correlated with total phosphorus concentrations (rho= 0.40). Drinking water for livestock was unaffected by trophic status, as was aquatic wildlife. The latter is not surprising given that the metric for their limitation is based on metals and the correlation between lead and total phosphorus, for example, was rather weak (rho=0.22).

Although trophic status appeared to qualify the relative ability of a river to provide these specific aquatic ecosystem services, we were surprised, as ecologists, that close to 50% of oligotrophic rivers were considered non-drinkable. Our a priori assumption was that oligotrophic ecosystems would provide all ecosystem services and that this would decline precipitously with increased eutrophication. Therefore, we tested how this assumption was supported by the data. To do so, we created an expected distribution of whether an ecosystem service was delivered or not by sequentially reattributing their expression per sampling event based on increasing trophic status. For example, in the case of irrigation, from the 6866 sampling events assessed, water was considered usable for irrigation 4663 times. These 4663 events were distributed among 2714 sites were oligotrophic, 2591 were mesotrophic and 1425 were eutrophic. Therefore 100% of the events in oligotrophic rivers are expected to be suitable for irrigation, based on our initial assumption, leaving 1949 drinkable events or 75% in the mesotrophic category, whereas none remained for the eutrophic category (expected percentage is 0%)(Figure 2c).

Using a Chi-squared test, we compared the observed versus the expected number of events for which each individual service could be provided (WebPanel 2). If trophic status is a perfect proxy of drinkability, as described above, we would expect that 90% of oligotrophic systems would be drinkable (Table S9); however, only 49% were considered drinkable (Table S10). In contrast, 0% of eutrophic events were expected to be drinkable, but we found that 20% were. Similar patterns were observed for recreation, irrigation, and the protection of aquatic wildlife, where fewer services were provided in oligotrophic ecosystems than expected whereas more were provided in eutrophic ones.

Therefore, trophic status is a good indicator of the general ability of a river to deliver ecosystem services because it serves as an indicator of a longer-term condition, but it is incomplete from a sampling event perspective, which may represent a more punctual situation. It is clear that trophic status co-varies much more with E. coli as compared to metals, but the incidence of E. coli compromising ecosystem services may be more sporadic in both space and time.

## 4. Conclusions

Here we show that using guideline thresholds for specific water usages is a novel integrative approach to characterize the capacity of a river to deliver on multiple aquatic ecosystem services. We found that classic ecological trophic status is a fair proxy to evaluate if water is safe for drinking, swimming, and irrigation. This is perhaps not too surprising since these ecosystem services are largely comprised by the concentration of E. coli where the land use changes that result in excess nutrient runoff from waste due to high human and/or animal populations are also subject to increasing incidence of E. coli. However, the frequency with which an ecosystem cannot provide an ecosystem service based on E. coli is sporadic. We suggest that trophic status represents the more overarching condition of an aquatic ecosystem, but that the inability to deliver on any ecosystem service based on specific guidelines is much more variable through time. For example, in terms of a more emergent contaminant of concern, the frequency that concentrations of microcystin, a cyanobacterial toxin, will limit the provisioning of different ecosystem services will be higher in more eutrophic waters, but this occurrence is never chronic (Taranu et al. 2017). The phenomenon with E. coli is similar in that concentrations are much more variable over time as compared to nutrients (Levy et al. 2009), but that the probability E. coli will exceed threshold guidelines is indeed higher in high-nutrient ecosystems. Therefore limiting excess nutrient runoff to surface water should remain a water quality maintenance priority.

Trophic status, however, did not serve as a good proxy for the suitability for livestock or the protection of aquatic wildlife given that their thresholds are limited by metal concentrations. Given the broad spatial extent of this study, elevated metal concentrations are most likely a function of natural geology or other types of anthropogenic activities (e.g. mining). However, the thresholds for the protection of wildlife are not a good indicator of whether an aquatic ecosystem is capable of maintaining its natural biodiversity. It is difficult to believe that 38% of all of oligotrophic events assessed were compromising biodiversity maintenance. Furthermore trophic status may be too simplistic as other anthropogenic pressures such as damming, over exploitation, and climate change are all contributing to habitat loss and aquatic biodiversity corrosion (Vorosmarty et al. 2010, Tonkin et al. 2019). Indeed given the global aquatic biodiversity crisis (IPBES 2018), there is an urgent need to manage aquatic ecosystems integratively and adaptively (Chan 2006). In order to meet the UN sustainable development goals of improving water quality and protecting life below water by 2030 (UN SDG 6 and 14 respectively), we believe that the novel use of thresholds to assess whether multiple ecosystems services will be of great use to managers and should be developed further. Indeed given the plethora of data collected routinely by government water quality surveys and different environmental programs, this information should be opened to help identify which ecosystems are most at risk.

## Acknowledgements

Financing for this project comes from NSERC Discovery to RM and JFL, OURANOS-Mitacs and a FQRNT fellowship to NFSG. This is contribution to the Strategic Network Lake Pulse and the GRIL.

## WebPanel1. Literature analysis

N. F. St-Gelais et al.

### 1.1 Temporal trends in the use of term water quality between 1980 and 2017 in ecology, biodiversity, agriculture, and public health

Using the Web of Knowledge Core Database, we downloaded metadata on all the scientific publications published between 1980 and 2017 with the term *water quality* in the title or abstract. In order to compare how the use of the term changed in ecological, public health, agricultural, and biodiversity sciences, we extracted metadata by field based on the Web of knowledge categories: *agronomy, agricultural eng., agriculture multi* for agriculture, *ecology* for ecology, *public, Environmental & occupational health* for public health and *biodiversity conservation* for biodiversity.

**Table S1:** Comparison of the segmented and linear model for each field. For the segmented regression each break point and the slope before and after each break point is reported.

| Fields | Slope-linear | AIC-linear | AIC-seg | Slope-seg-1 | BreakPoint1 | Slope-seg-2 | BreakPoint2 | Slope-seg-3 |
| --- | --- | --- | --- | --- | --- | --- | --- | --- |
| General | 0.09 | -24 | -33 | 0.00 | 1985 | 0.09 |  |  |
| Agriculture | 0.08 | 59 | 36 | 0.01 | 1987 | 0.15 | 2002 | -0.01 |
| Ecology | 0.09 | 11 | -3 | 0.00 | 1987 | 0.11 | 2008 | 0.03 |
| Public Health | 0.09 | -24 | -33 | 0.00 | 1985 | 0.09 |  |  |
| Biodiversity | 0.08 | 45 | 21 | 0.04 | 2001 | 0.15 |  |  |

To compare temporal trends in the use of the term *water quality* relative to the total number of publications in each field, we used a z-score approach. For each field, we fitted both a linear and segmented regression (*segmented* R package) model, and selected the best model fit to the data using the Akaike information criterion (AIC). The relative use of the term *water quality* increased linearly in the overall literature (Table S1). However the segmented model often better fit the observed data when we considered each field separately. Indeed, for agriculture and ecology, we observed a sharp increase starting in the late eighties followed by a stabilization in the use of the term around 2010. In biodiversity science, we observed a similar increase in the late eighties followed by an acceleration that started in 2001 that continues to increase until 2017. For public health, we observed a break point around 1985. However, this breakpoint was not considered significant because it only captured the high initial variability caused by the very small number of studies that included the term *water quality*, so the linear model was selected.

**Table S2:** The 20 most frequently used keywords in each Web of knowledge categories

| Rank | Ecology | Agriculture | Public Health | Biodiversity and conservation |
| --- | --- | --- | --- | --- |
| 1 | land | management | Health | environment |
| 2 | ecosystem | soil | Source | land |
| 3 | management | land | Drinking | monitoring |
| 4 | environmental | source | Risk | species |
| 5 | river | nitrogen | contamination | river |
| 6 | nutrient | crop | Coli | environmental |
| 7 | species | field | Treatment | ecosystem |
| 8 | time | treatment | Public | management |
| 9 | phosphorus | surface | concentrations | indicators |
| 10 | stream | flow | population | assessment |
| 11 | increased | agricultural | Standards | region |
| 12 | surface | production | Control | aquatic |
| 13 | scale | nutrient | Human | ecological |
| 14 | ecological | parameters | environmental | habitat |
| 15 | concentrations | increased | parameters | urban |
| 16 | nitrogen | control | household | ecosystems |
| 17 | flow | phosphorus | Local | local |
| 18 | soil | time | community | concentrations |
| 19 | lake | nitrate | Surface | parameters |
| 20 | spatial | concentrations | intervention | status |
| 21 | sediment | practices | Pollution | abundance |
| 22 | aquatic | environmental | Rural | index |
| 23 | habitat | irrigation | Urban | diversity |
| 24 | impacts | impacts | waterborne | lake |
| 25 | treatment | values | management | increased |

### 1.2 Comparing the words most often associated with water quality among fields in 2017

To further explore how the focus in the use of the term *water quality* differs among fields we extracted the words most often associated with *water quality* in the title and abstracts. The titles and abstracts from manuscripts published in 2017 only in each of the four fields were first downloaded from Web of Knowledge. After removing words that were not meaningful, we calculated the relative frequency of each word. To quantitatively compare how the words associated with *water quality* differed between the different fields, we measured the cosine distance (from the *lsaR* R package) to assess how different the top 100 words were (Table S3).

**Table S3:** Cosine distance between the words associated to *water quality* in each field.

| Ecology | Biodiversity | Public Health | Agriculture |
| --- | --- | --- | --- |
| 1.00 | 0.71 | 0.17 | 0.55 |
| 0.71 | 1.00 | 0.31 | 0.36 |
| 0.17 | 0.31 | 1.00 | 0.15 |
| 0.55 | 0.36 | 0.15 | 1.00 |

## WebPanel2. Ecosystem services in Canadian Rivers

N. F. St-Gelais et al.

### 2.1 Aggregated guideline database

We created an aggregated guideline database using Canadian guidelines published by the Canadian Council of Ministers of the Environment (CCME) and Health Canada for the following ecosystem services: drinking, swimming, irrigation, livestock, and the protection of aquatic wildlife. We only included guidelines with a quantitative numeric threshold and as such, narrative guidelines were excluded. Furthermore, using guidelines that are under the detection limit (e.g. dicamba for irrigation) would result in artificially limiting the service each time it was measured. In those rare cases, we used the detection limit multiplied by 2 as a modified guideline which we specified in the database. We also included trophic status guidelines from Dodds 1998 classification for rivers.

We consolidated river water quality data from the two following openly available data sources: the National Long-term Water Quality Monitoring Data and the Provincial (stream) water quality monitoring network (PWQMN) from Ontario, for a total of 46 databases.

**Table S4:**
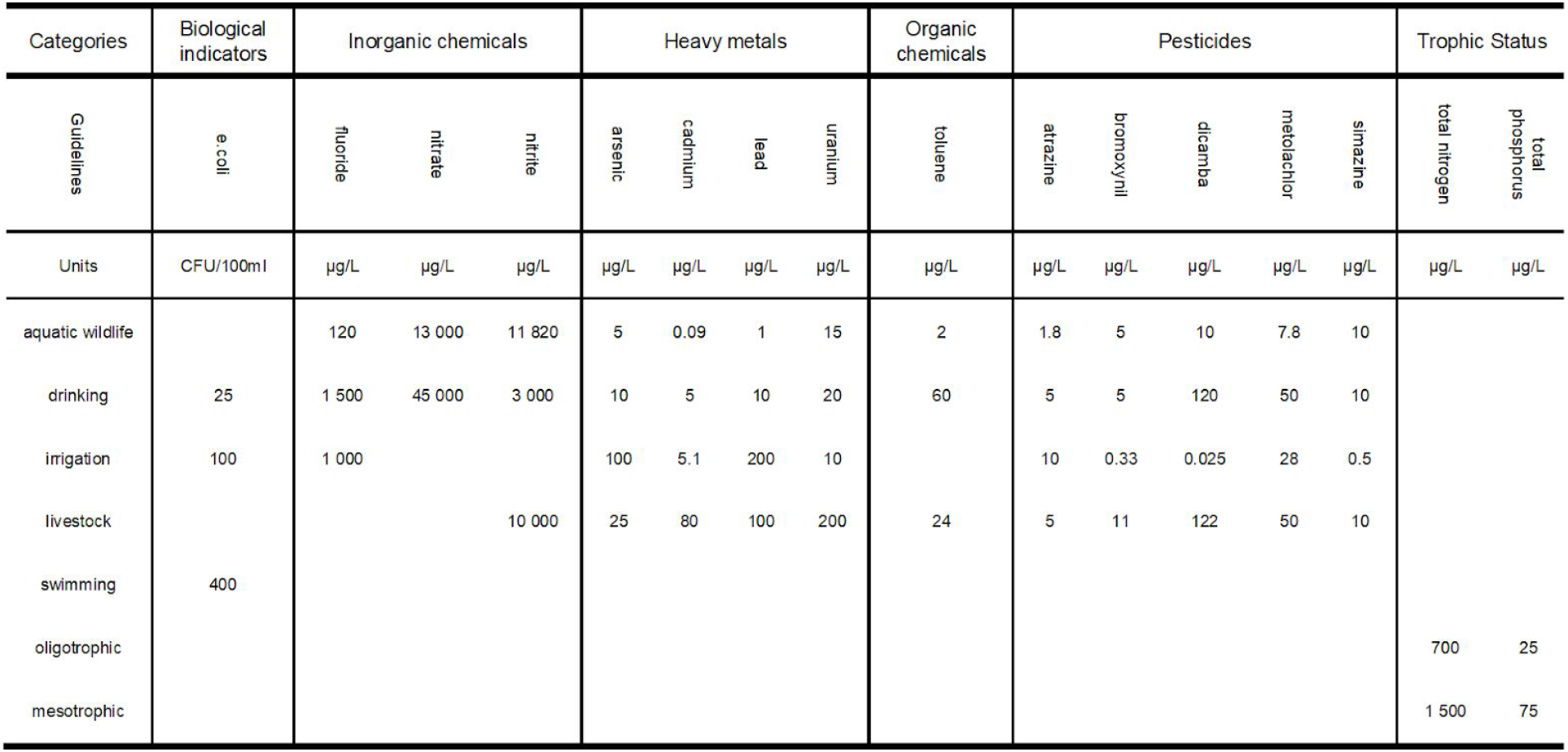
Table on selected guidelines.

### 2.2 Aggregated limnological database

We developed the *dbExtract* R package and ETL (extract, transform and load) package to 1) combine and normalize the databases, 2) evaluate which ecosystem service(s) can potentially by provided at each sampling event, and 3) evaluate trophic status based on the Dodds 1998 classification. In the combined database, we had a total of 660 000 unique observations from rivers across Canada over 22 years (from 1995 to 2017). We grouped observations by station and by month to create an aggregated limnological database of observations for a total of 65935 sampling events. As there are multiple guidelines for each ecosystem service (e.g. close to 75 for drinking water), for this study we selected a subset of guidelines in each group (Table S4) based on their measurement frequency in publicly available databases as well as for their use in assessing multiple ecosystem services.

**Table S5:** Cosine distances between ecosystem services based on the selected guidelines

| ES | aquatic wildlife | drinking | irrigation | livestock | swimming | mesotrophic | oligotrophic |
| --- | --- | --- | --- | --- | --- | --- | --- |
| aquatic wildlife | 1.00 | 0.96 | 0.84 | 0.92 | 0.00 | 0.00 | 0.00 |
| drinking | 0.96 | 1.00 | 0.89 | 0.89 | 0.27 | 0.00 | 0.00 |
| irrigation | 0.84 | 0.89 | 1.00 | 0.82 | 0.30 | 0.00 | 0.00 |
| livestock | 0.92 | 0.89 | 0.82 | 1.00 | 0.00 | 0.00 | 0.00 |
| swimming | 0.00 | 0.27 | 0.30 | 0.00 | 1.00 | 0.00 | 0.00 |
| mesotrophic | 0.00 | 0.00 | 0.00 | 0.00 | 0.00 | 1.00 | 1.00 |
| oligotrophic | 0.00 | 0.00 | 0.00 | 0.00 | 0.00 | 1.00 | 1.00 |

We developed the *load_ES* function to evaluate, for each sampling event in the aggregated limnological DB, which ecosystem services could potentially be provided or not based on the guideline threshold. Because there was considerable variability in which variables were measured at each sampling event, this evaluation was based on the guideline for which specific information was available. In the case when no variable was available to evaluate whether a service could be provided or not, the service wasn’t evaluated for this specific sampling event.

### 2.3 Calculating limiting frequency for each guidelines

For each of the selected guidelines (see section 2.1), we assessed the frequency with which each was over the threshold as a function to the total times it was measured. As such, for each sampling event multiple guidelines could be limiting an ecosystem service, with the exception of swimming which only had E. coli concentration as a guideline metric (see Table S4).

We observed that in average E. coli was often limiting the following ecosystem services: swimming, irrigation, and drinking (see section 2.3 and Table S6), but was only measured in 11% of sampling events (see Table S6). Hence, because our approach is based on the variables measured during each sampling event, this could lead to an overestimation of how often an ES is provided, particularly for ecosystem services that had more than one guideline such as irrigation and drinking. To avoid this bias, for the analyses we used a subset of sampling events for which E. coli was measured.

**Table S6:**
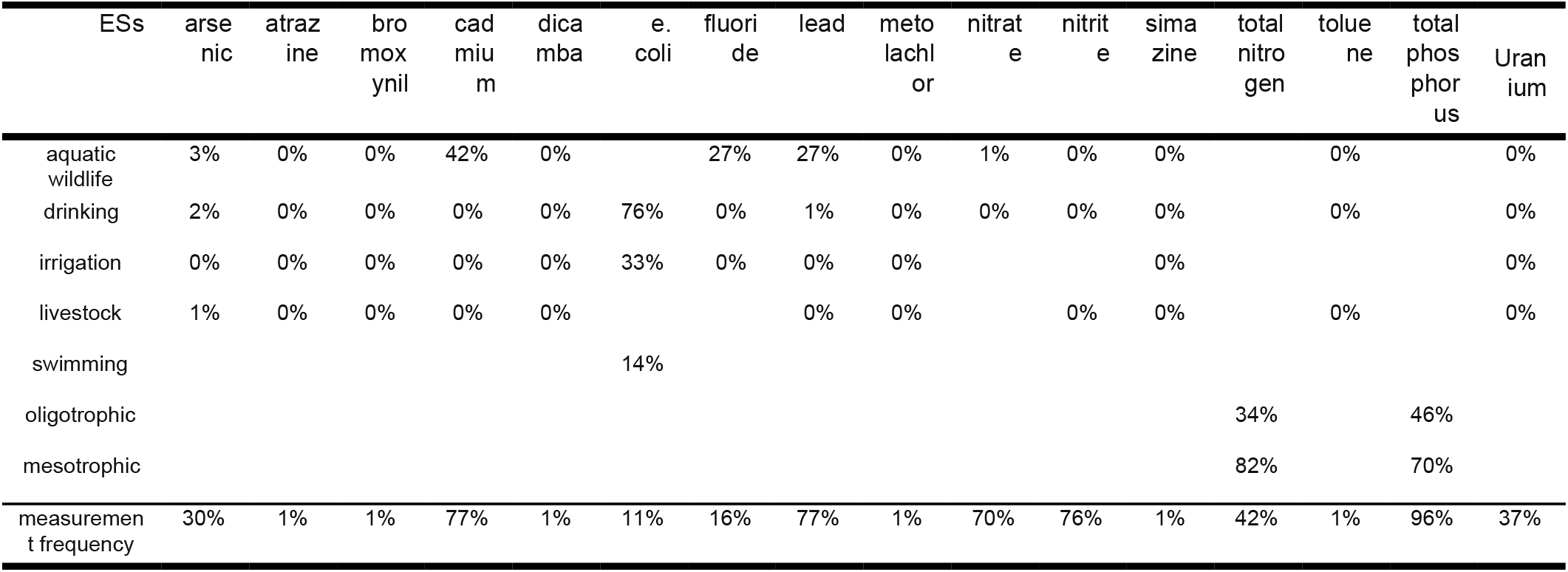
Guidelines limiting frequency in percentage. The measurement frequency of each compound is also reported.

**Table S7:**
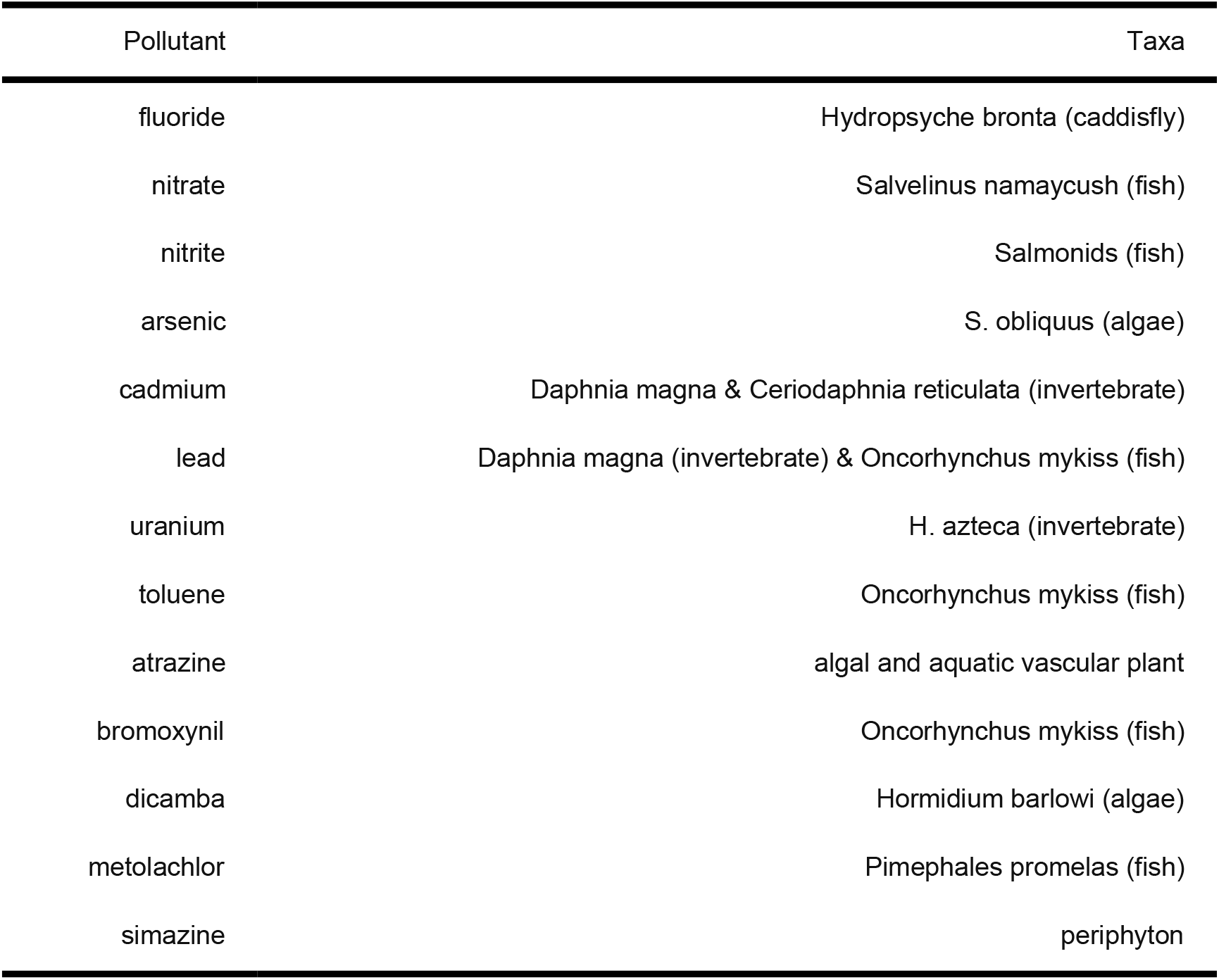
Most sensitive taxonomic group(s) on which the long-term aquatic wildlife protection guideline is based.

|  |  |
| --- | --- |
| fluoride | Hydropsyche bronta (caddisfly) |
| nitrate | Salvelinus namaycush (fish) |
| nitrite | Salmonids (fish) |
| arsenic | S. obliquus (algae) |
| cadmium | Daphnia magna & Ceriodaphnia reticulata (invertebrate) |
| lead | Daphnia magna (invertebrate) & Oncorhynchus mykiss (fish) |
| uranium | H. azteca (invertebrate) |
| toluene | Oncorhynchus mykiss (fish) |
| atrazine | algal and aquatic vascular plant |
| bromoxynil | Oncorhynchus mykiss (fish) |
| dicamba | Hormidium barlowi (algae) |
| metolachlor | Pimephales promelas (fish) |
| simazine | periphyton |

### 2.4 Testing if oligotrophic rivers are providing more ecosystem services: observed versus expected

In order to verify our hypothesis that more ecosystem services are provided in oligotrophic than in mesotrophic and eutrophic rivers, we statistically tested, using a chi-squared test, if the distribution of events by trophic state is in agreement with the hypothesis that trophic status is a perfect proxy for whether or not an ecosystem service can be provided or not. To test this, we did the following:

1. For each ecosystem service, we selected sites for which the ecosystem service and trophic state could be evaluated: at least one guideline was measured as well as total phosphorus or total nitrogen.
2. We created the observed values table (Table S9) by calculating the number of sampling events for which an ecosystem service could be provided as a function of true trophic status.
3. To create the expected table (Table S10), we attributed all the sampling events for which an ecosystem service could be provided in order of oligotrophic, then mesotrophic and finally eutrophic sampling events. In other words, by default we attributed all the events for which a service could be provided to oligotrophic systems, and subsequently to mesotrophic systems and in the rare case to eutrophic rivers. Then we plotted the observed and expected in a radar plot based frequency of delivery as a function of trophic status.

For clarity we repeat the example as stated in the paper: in the case of irrigation, from the 6866 sampling events assessed, water was considered usable for irrigation 4663 times. These 4663 events were distributed among 2714 sites were oligotrophic, 2591 were mesotrophic and 1425 were eutrophic. Therefore 100% of the events in oligotrophic rivers are expected to be suitable for irrigation, based on our initial assumption, leaving 1949 drinkable events or 75% in the mesotrophic category, whereas none remained for the eutrophic category.

**Table S8:** Chi-squared table. Observed (obs) and expected (exp) are reported in percentages. The Pearson residuals (res) represent the relative contribution of each trophic state to the chi-square statistic, where larger residuals indicate a greater deviance from the expected distribution.

| Ecosystem services | oligo.obs | oligo.exp | oligo.res | meso.obs | meso.exp | meso.res | eutro.obs | eutro.exp | eutro.res | p-value |
| --- | --- | --- | --- | --- | --- | --- | --- | --- | --- | --- |
| aquatic | 0.53 | 1.00 | -24.93 | 0.38 | 0.03 | 95.81 | 0.27 | 0.00 | 395.00 | 0.00 |
| drink | 0.49 | 0.90 | -22.51 | 0.31 | 0.00 | 831.00 | 0.20 | 0.00 | 299.00 | 0.00 |
| irrigation | 0.85 | 1.00 | -8.06 | 0.66 | 0.75 | -5.62 | 0.47 | 0.00 | 668.00 | 0.00 |
| livestock | 1.00 | 1.00 | 0.00 | 1.00 | 1.00 | -0.06 | 1.00 | 1.00 | 0.08 | 1.00 |
| recreational | 0.96 | 1.00 | -1.90 | 0.85 | 1.00 | -6.73 | 0.61 | 0.29 | 21.24 | 0.00 |

**Table S9:**
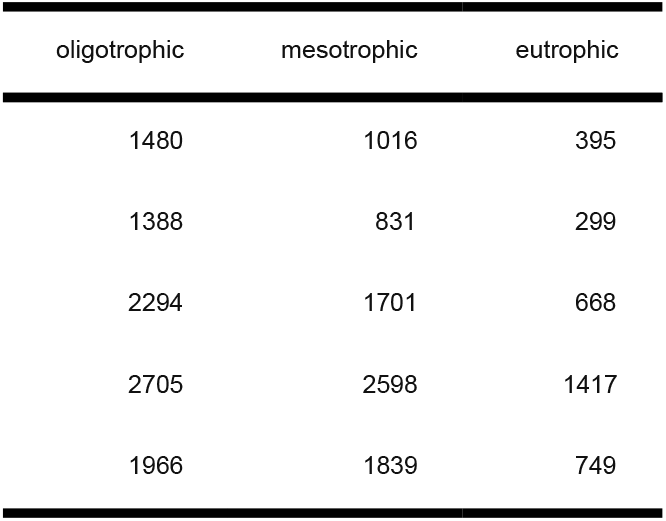
Observed distribution of sampling events for which each ecosystem service could be provided by trophic state.

**Table S10:** Expected distribution of sampling events for which each ecosystem service could be provided by trophic state.

| oligotrophic | mesotrophic | eutrophic |
| --- | --- | --- |
| 2799 | 92 | 0 |
| 2518 | 0 | 0 |
| 2714 | 1949 | 0 |
| 2705 | 2601 | 1414 |

## References

Anders, P. J. and Ashley, K. I. 2007. The Clear-water Paradox of Aquatic Ecosystem Restoration. - Fisheries 32: 125–128.

Boyd, D. R. 2006. The water we drink: an international comparison of drinking water quality standards and guidelines.

Canadian Council of Ministers of the Environment 1999. Canadian environmental quality guidelines.

Canadian Council of Ministers of the Environment 2003. Canadian water quality guidelines for the protection of aquatic life: Guidance on the Site-Specific Application of Water Quality Guidelines in Canada: Procedures for Deriving Numerical Water Quality Objectives. - In: Canadian environmental quality guidelines.

Canadian Environmental Protection Act (CEPA). 1988.

Canada Fisheries Act. R.S.C., 1985, c. F-14. 1985.

Chan, K. M. A. et al. 2006. Conservation Planning for Ecosystem Services. - PLOS Biology 4: e379.

Daily, G. C. et al. 2009. Ecosystem services in decision-making: time to deliver. - Frontiers in Ecology and the Environment 7: 21–28.

Dodds, W. 1998. Suggested classification of stream trophic state: distributions of temperate stream types by chlorophyll, total nitrogen, and phosphorus - ScienceDirect. - Water Research 32: 1455–1462.

Dodds, W. K. 2006. Eutrophication and trophic state in rivers and streams. - Limnol. Oceanogr. 51: 671–680.

Gordon, L. J. et al. 2010. Managing water in agriculture for food production and other ecosystem services. - Agricultural Water Management 97: 512–519.Beni

Guerry, A. D. et al. 2015. Natural capital and ecosystem services informing decisions: From promise to practice. - PNAS 112: 7348–7355.

Hart, B. T. et al. 1993. Australian water quality guidelines: a new approach for protecting ecosystem health. - J Aquat Ecosyst Stress Recov 2: 151–163.

Health Canada 2012. Guidelines for Canadian Recreational Water Quality, Third Edition. - Water, Air and Climate Change Bureau, Healthy Environments and Consumer Safety Branch, Health Canada.

Health Canada 2017. Guidelines for Canadian Drinking Water Quality.

IPBES 2018. The IPBES regional assessment report on biodiversity and ecosystem services for the Americas.: 656.

Levy, K. et al. 2009. Drivers of Water Quality Variability in Northern Coastal Ecuador. - Environ. Sci. Technol. 43: 1788–1797.

Mace, G. M. et al. 2012. Biodiversity and ecosystem services: A multilayered relationship. - Trends in Ecology and Evolution 27: 19–25.

Mateo-Sagasta, J. et al. 2017. Water pollution from agriculture: a global review.

Orth, R. J. et al. 2006. A Global Crisis for Seagrass Ecosystems. - BioScience 56: 987–996.

Raudsepp-Hearne, C. et al. 2010. Ecosystem service bundles for analyzing tradeoffs in diverse landscapes. - Proceedings of the National Academy of Sciences 107: 5242–5247.

Smith, V. H. 2003. Eutrophication of freshwater and coastal marine ecosystems a global problem. - Environ Sci & Pollut Res 10: 126–139.

Taranu, Z. E. et al. 2017. Predicting microcystin concentrations in lakes and reservoirs at a continental scale: A new framework for modelling an important health risk factor. - Global Ecol. Biogeogr. 26: 625–637.

Tonkin, J. D. et al. 2019. Prepare river ecosystems for an uncertain future. - Nature 570: 301.

UNEP 2016. A Snapshot of the World’s Water Quality: Towards a global assessment.: 162.

U.S. Water Quality Act of 1987. Pub.L. 100-4, February 4, 1987.

UK Water Act. c. 15. 1989.

Vörösmarty, C. J. et al. 2010. Global threats to human water security and river biodiversity. - Nature 467: 555–561.

Ziegler, J. P. et al. 2017. Social-ecological outcomes in recreational fisheries: the interaction of lakeshore development and stocking. - Ecological Applications 27: 56–65.

